# Increasing the Efficiency of Genome-wide Association Mapping via Hidden Markov Models

**DOI:** 10.1101/039099

**Authors:** Hong Gao, Hua Tang, Carlos D. Bustamante

## Abstract

With the rapid production of high dimensional genetic data, one major challenge in genome-wide association studies is to develop effective and efficient statistical tools to resolve the low power problem of detecting causal SNPs with low to moderate susceptibility, whose effects are often obscured by substantial background noises. Here we present a novel method that serves as an optimal technique for reducing background noises and improving detection power in genome-wide association studies. The approach uses hidden Markov model and its derivate Markov hidden Markov model to estimate the posterior probabilities of a markers being in an associated state. We conducted extensive simulations based on the human whole genome genotype data from the GlaxoSmithKline-POPRES project to calibrate the sensitivity and specificity of our method and compared with many popular approaches for detecting positive signals including the *χ*^2^ test for association and the Cochran-Armitage trend test. Our simulation results suggested that at very low false positive rates (< 10^−6^), our method reaches the power of 0.9, and is more powerful than any other approaches, when the allelic effect of the causal variant is non-additive or unknown. Application of our method to the data set generated by Welcome Trust Case Control Consortium using 14,000 cases and 3,000 controls confirmed its powerfulness and efficiency under the context of the large-scale genome-wide association studies.

## Introduction

The International HapMap project [1-3] and associated advances in large-scale genotyping technology have enabled genome-wide association studies (GWAS) of common genetic variants involved in major human diseases [4-6]. For example, the pioneering studies by Abecasis et al. [7] and Klein et al. [8] have demonstrated the effectiveness and powerfulness of the GWAS for finding susceptible loci associated with the multifactorial disease age-related macular degeneration. Likewise, Wellcome Trust Case-Control Consortium [6] identified 24 independent genetic variants that potentially contribute to seven complex human diseases through large-scale whole genome association studies using 14,000 cases and 3,000 controls, genotyped across 500,000 SNPs. Since then, numerous GWAS projects discovered common genetic variants for a wide variety of human diseases, signifying that the era of clinical genomics has arrived.

The technology enabling GWAS has also created a critical need for novel statistical tools for analyzing the copious amounts of data generated by these projects (e.g., millions of SNPs queried for association with disease phenotypes across thousands of individuals). A particular pressing problem is that in most GWAS studies, the low-power single marker analysis methods, e.g., the *χ*^2^ test for allelic or genotype association and the Cochran-Armitage trend test [9], are used most often in detecting susceptible genetic variants. Along with single marker analysis, comes the multiple comparison problem, which arises when individually testing each SNP for association with disease phenotype. Many investigators choose to address this problem by performing Bonferroni correction [8, 10], even though it is notorious for being overly conservative. Other popular approaches for solving this problem include controlling for false discovery rate (FDR) or positive FDR for its variants. However, because these approaches assume independence among tests, they do not work well for association studies [11-14].

Another way to address the problem is to look at it from a different perspective. Human genomes can be viewed as an assembly of consecutive blocks of strong linkage disequilibrium (LD), within which multiple SNPs are closely correlated with each other [4]. For any given association study, only a very small number of blocks contain risk SNPs or haplotypes related to the disease. Several approaches have been proposed to use hidden Markov model (HMM) to predict the locations of such blocks across genomes. The HMM paradigm has been particularly successful in admixture mapping, where a few thousand SNPs are used to tag chromosomal segments from ancestral populations [15-17]. However, the Markovian assumption of HMM appears to be invalid among a dense set of markers due to “background linkage disequilibrium” [18], thus cautions need to be taken in proper applications of HMM to dense association studies. Recognizing this shortcoming of HMM, Tang et al. [19] introduced additional dependencies among neighboring markers into HMM and developed the Markov-Hidden Markov Model (MHMM). This method inherits the computationally efficient and succinct framework from HMM but also takes into account correlation among markers.

In this paper, we develop novel HMM and MHMM methods for GWAS. Our approach to fine-scale mapping makes efficient use of the correlation among markers to strike an optimal balance between power and Type I error. Informaly, our approach use the Hidden Markov Model or the Markov Hidden Markov Model to search for segments of sequences showing significant association signals. This HMM / MHMM approach has little computational overhead compared to single marker analysis, and statistically delineates potential susceptibility regions of the genome. Furthermore, our method has several advantages over other approaches. For example, it depends on only a few assumptions about the underlying genetic model of disease and provides a flexible framework into which various genetic factors can be easily incorporated. Most importantly, it naturally avoids the multiple comparison problem and its power increases with sample size and the number of markers, a desirable property of any approach for genome-wide association mapping.

## Results

We developed HMM and MHMM based methods to utilize the correlation information among markers to achieve fine-scale disease mapping. Here the trait values of individuals can be either categorical (e.g. in case-control studies) or quantitative. In the data application, we focus on qualitative phenotypes, especially the case-control context. Both the Hidden Markov Model and the Markov Hidden Markov Model assume two hidden states, the associated state and the not-associated state of the marker with a phenotype. The three major components of the HMM / MHMM frameworks, including the prior distribution of the two states, the transition probability matrix, and the emission probabilities, are specified in Methods Section.

### Application to Simulated Data

To test the accuracy and robustness of our methods in GWAS applications, we used the European genotype data from GlaxoSmithKline (GSK)-POPRES project [20] to perform simulation in order to retain the true LD pattern of human genome, as illustrated in Figure 1. Due to the pure composition of our dataset, the LD extends longer than other studies across distinct human groups [41]. 100 replications were generated based on this data set using the multiplicative risk model for each of the four penetrance sets with the relative risk (RR) ranging from 1.25 to 2 assuming either additive or recessive allelic effect (see Table 1). Detailed simulation scheme is illustrated in Methods Section. We applied our HMM and MHMM methods to the simulated data assuming the prior probability of associated state *β* = 10^−6^, the LD measure λ = 1/15*kb*, and the non-centrality parameter *κ* = 30.0 based on empirical information. Our method is strikingly efficient - it takes only a couple of minutes to run on a data set with 500,000 markers and 1,000 individuals. Its short running time and ability to handle high dimensional data makes it a very powerful tool for genome-wide studies. Since we placed such a high value on efficiency, we would prefer not to use the MCMC algorithm, which brings with it extra problems like convergence issue.

**Figure 1.**
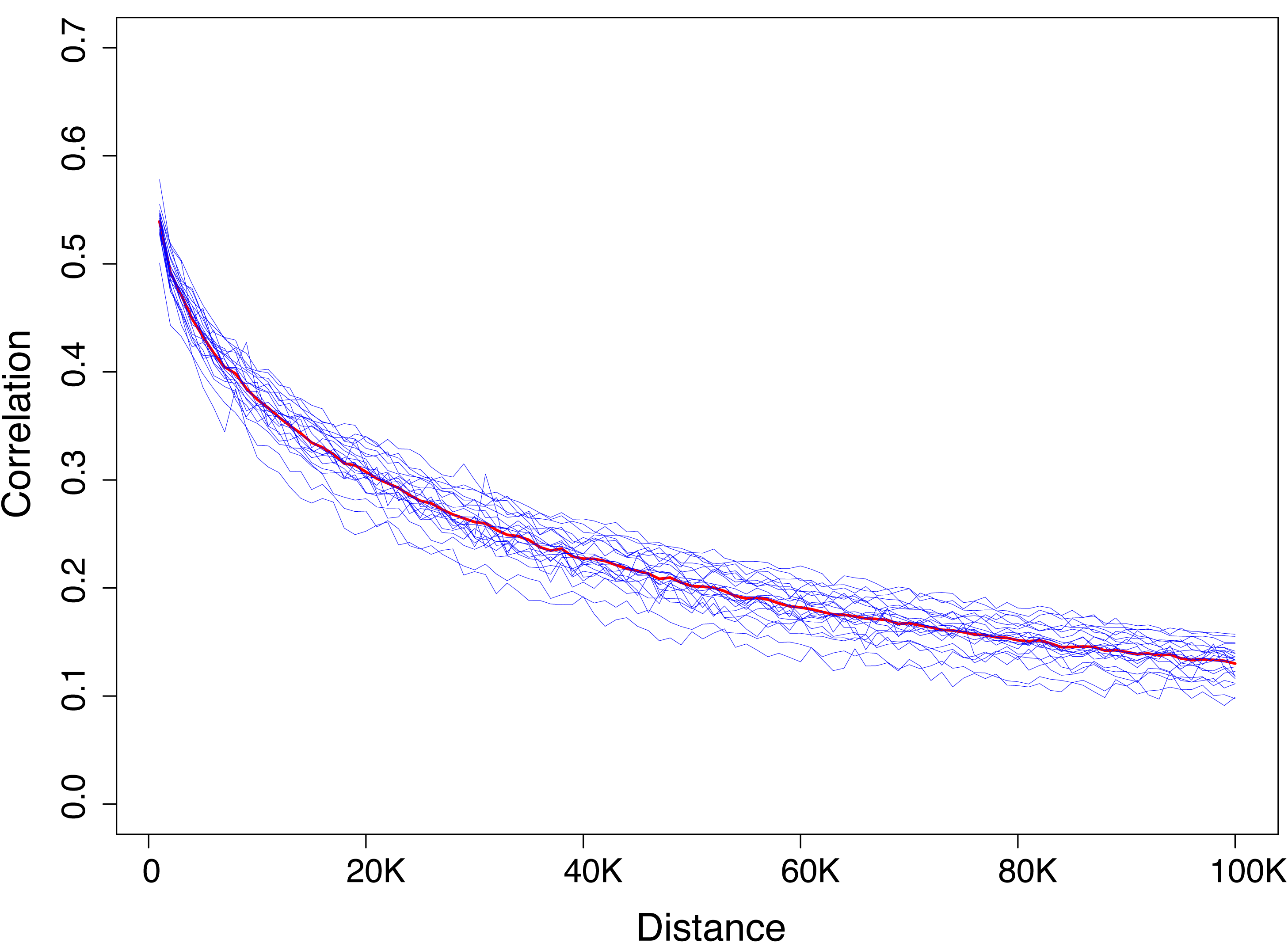
The LD decay pattern based on 22 autosomes of selected 1000 Europeans. Each blue line represents the decrease of LD measure, the correlation coefficient *r*^2^, as a function of distance among two markers along one of 22 autosomes. The red line is an average of the 22 blue lines, i.e., averaged LD decay pattern across genomes.

**Table 1.** Penetrance sets for simulation

| Penetrance Set | $\mathbb{P}(D \mid AA)$ | $\mathbb{P}(D \mid Aa)$ | $\mathbb{P}(D \mid aa)$ | RR |
| --- | --- | --- | --- | --- |
| Additive I | 0.5 | 0.45 | 0.4 | 1.25 |
| Additive II | 0.6 | 0.5 | 0.4 | 1.5 |
| Additive III | 0.7 | 0.55 | 0.4 | 1.75 |
| Additive IV | 0.8 | 0.6 | 0.4 | 2 |
| Recessive I | 0.5 | 0.4 | 0.4 | 1.25 |
| Recessive II | 0.6 | 0.4 | 0.4 | 1.5 |
| Recessive III | 0.7 | 0.4 | 0.4 | 1.75 |
| Recessive IV | 0.8 | 0.4 | 0.4 | 2 |

Four penetrance sets are used in the simulation. "D" represents disease. "AA", "Aa" and "aa" represent possible genotypes occurring at each SNP locus. "RR" represent the relative risk of the locus, a measure of susceptibility of a disease, equal to $\frac{\mathbb{P}(D|AA)}{\mathbb{P}(D|aa)}$ .

We assessed the performance of this novel method in comparison with the *χ*^2^ test for genotypic association and the Cochran-Armitage trend test by drawing the receiver operating characteristic (ROC) curves for each method [21,22]. ROC curves depict relative trade-offs between true positive and false positive. We define a true positive based on distance from the causal SNP using either a 20kb or 100kb threshold. Likewise, associated markers beyond this threshold were classified as false positive signals. We defined true positive rates per penetrance set as the percentage of true positives out of 100, which was the total number of replications considered. The false positive rate was defined as the proportion of negatives that were false positives.

For additive penetrance set I, due to the substantially low relative risk (*RR* = 1.25), the effects of the susceptible alleles are close to the background signals and are difficult to detect. As illustrated in Figure 2A, when the power of HMM / MHMM reaches 0.5, their Type I error rate is controlled at 10^−3^ scale. At the same level of power, the HMM and MHMM methods have lower false positive error rate than either the *χ*^2^ genotypic association test or the Cochran-Armitage trend test. This indicates our HMM/MHMM approach is an effective tool in eliminating the effects of background noises, especially at this low susceptibility level. However, when the true signal is so weak and hard to distinguish from background signals, our methods sometimes obscure the true signal along with background noises. That’s the reason why we cannot obtain 100% power even when the false positive rate increases to a very large value. The MHMM and the HMM are similarly accurate, as their ROC curves almost overlap each other. When the distance threshold from the causal SNP is reduced from 100 KB to 20 KB, the ROC curves shift to the direction of higher false positive rates. This implies that it is still impractical in most cases to fine map a low susceptible disease locus with only 1000 individuals.

**Figure 2.**
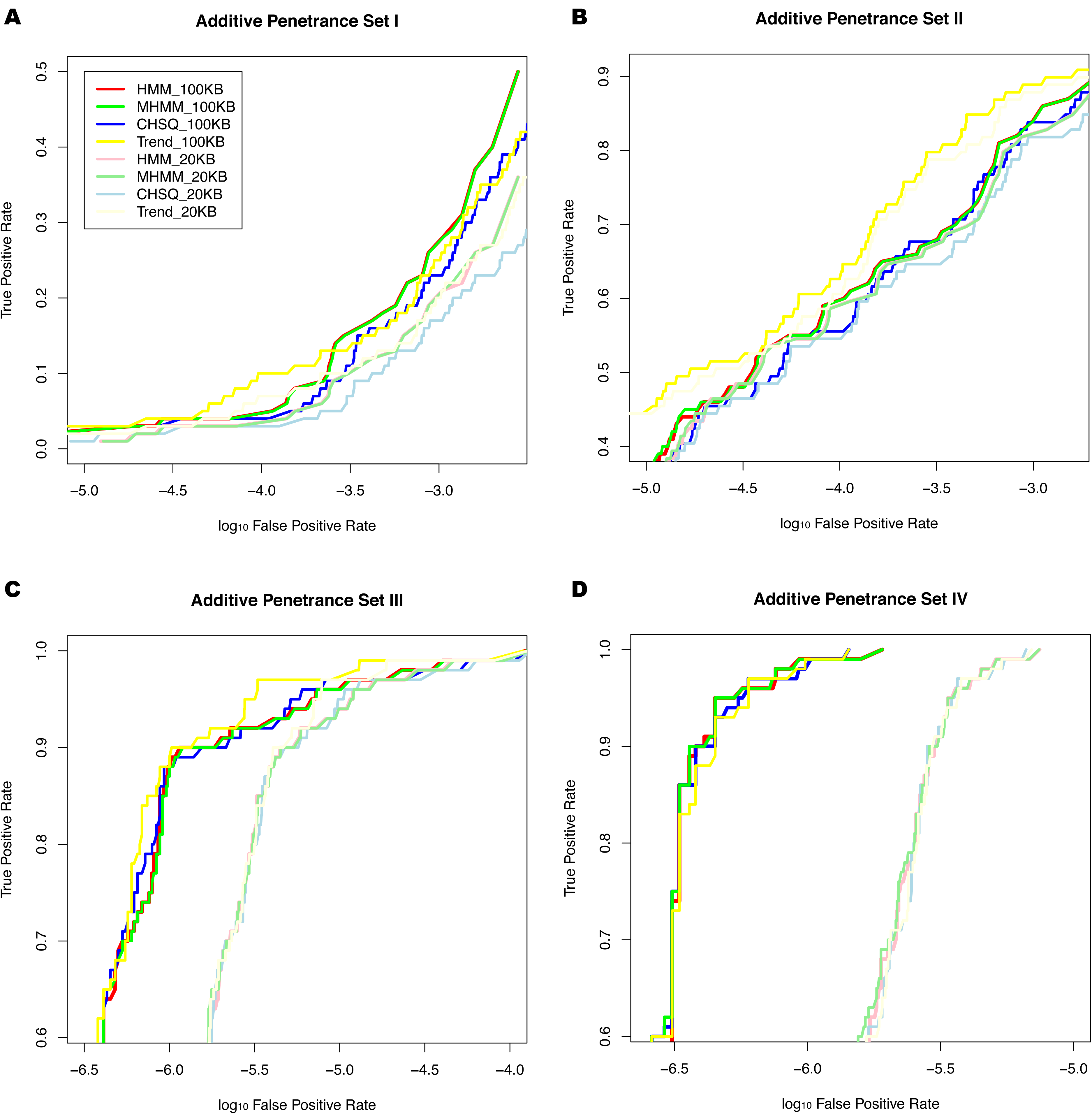
ROC curves of HMM / MHMM methods, *χ*^2^ genotypic association test, and Cochran-Armitage trend test for four additive penetrance sets. A, B, C, and D are plots for additive penetrance set I, II, III, and IV, respectively. The red lines represent the ROC curves of HMM method; the green lines are for MHMM; blue for *χ*^2^ genotypic association test; yellow for Cochran-Armitage trend test. “100KB” or “20KB” means the distance threshold from the causal variant is 100 kb or 20kb, represented by the normal and light colors, respectively.

When the true signal’s relative risk increases to 1.5, the HMM / MHMM approaches grow much more accurate, shown in Figure 2B. Their power increase to nearly 0.6, when the false positive rates are controlled at 10^−4^ levels, and at the 0.4 level of power, the Type I error rates decrease to 10^−5^, an acceptable level for genome-wide association studies. The performance of the MHMM mostly overlaps that of the HMM approach, and both are generally better than the *χ*^2^ genotypic association test. When the distance threshold from the causal SNP region decreases from 100 KB to 20 KB, the power no longer decreases so obviously, implying fine mapping becomes practical when the relative risk of the disease exceeds a detection threshold. The Cochran-Armitage trend test outperforms the rest three methods in this case. That’s probably due to the fact that it is locally most powerful when the allelic effect is exactly additive.

In the additive penetrance set III, the relative risk is increased to 1.75 and the false positive rates of the HMM and the MHMM are well controlled at 10^−6^ scales even when their power increase to 0.8 (see Figure 2C). When the power increases to close to 1, they still have very low false positive rates around 10^−5^. The MHMM generally has the same pattern as the HMM, and the performance of the *χ*^2^ genotypic association test is very similar to that of HMM / MHMM. The Cochran-Armitage trend test still performs better than the other three methods. When the distance cutoff from the true polymorphism decreases from 100 KB to 20 KB, the false positive rates inflate almost one magnitude. This might be due to the fact that when the signal is dramatically strong and according to Figure 1 the LD of the data set extends beyond 100KB on average, markers nearby or within a reasonable long distance are still in tight correlation with it. Therefore, the narrow covering region might inflate the false positive rate.

When the relative risk of the causal variant increases to a considerably large value like 2.0, all the methods have no problem detecting true positive signals at a very low level of Type I error rates (see Figure 2D). In this case, both the HMM and MHMM have nearly perfect power at the false positive level of 10^−6^. At this large value of relative risk, fine mapping becomes feasible because the power for the 20 KB distance threshold from the true polymorphism is close to 1 at the false positive level of 10^−5.5^. The power of the *χ*^2^ test also reaches a high level, but it still underperforms the HMM and MHMM slightly at a few false positive levels. The Cochran-Armitage trend test performs slightly worse than the other three methods. These results indicate that when the susceptibility of the disease-related SNP is high enough, most association mapping approaches can detect the effect.

Considering the recessive allelic effect, the patterns of ROC curves (shown in Figure 3) for four methods are largely similar to that under additive allelic effect across different values of relative risk. The major difference is that the Cochran-Armitage trend test performs much worse than the other three methods for different levels of relative risk, as its additive assumption is not satisfied in this recessive case. The HMM and the MHMM generally outperform or at least perform as well as the *χ*^2^ test. When the relative risk of the causal variant is large, such as 1.75 or 2.0, the MHMM outperforms the HMM, which demonstrates the advantage of correcting for the background LD. When the distance threshold from the causal variant shrinks to 20 KB, the false positive rates inflate almost one magnitude for various levels of relative risk.

**Figure 3.**
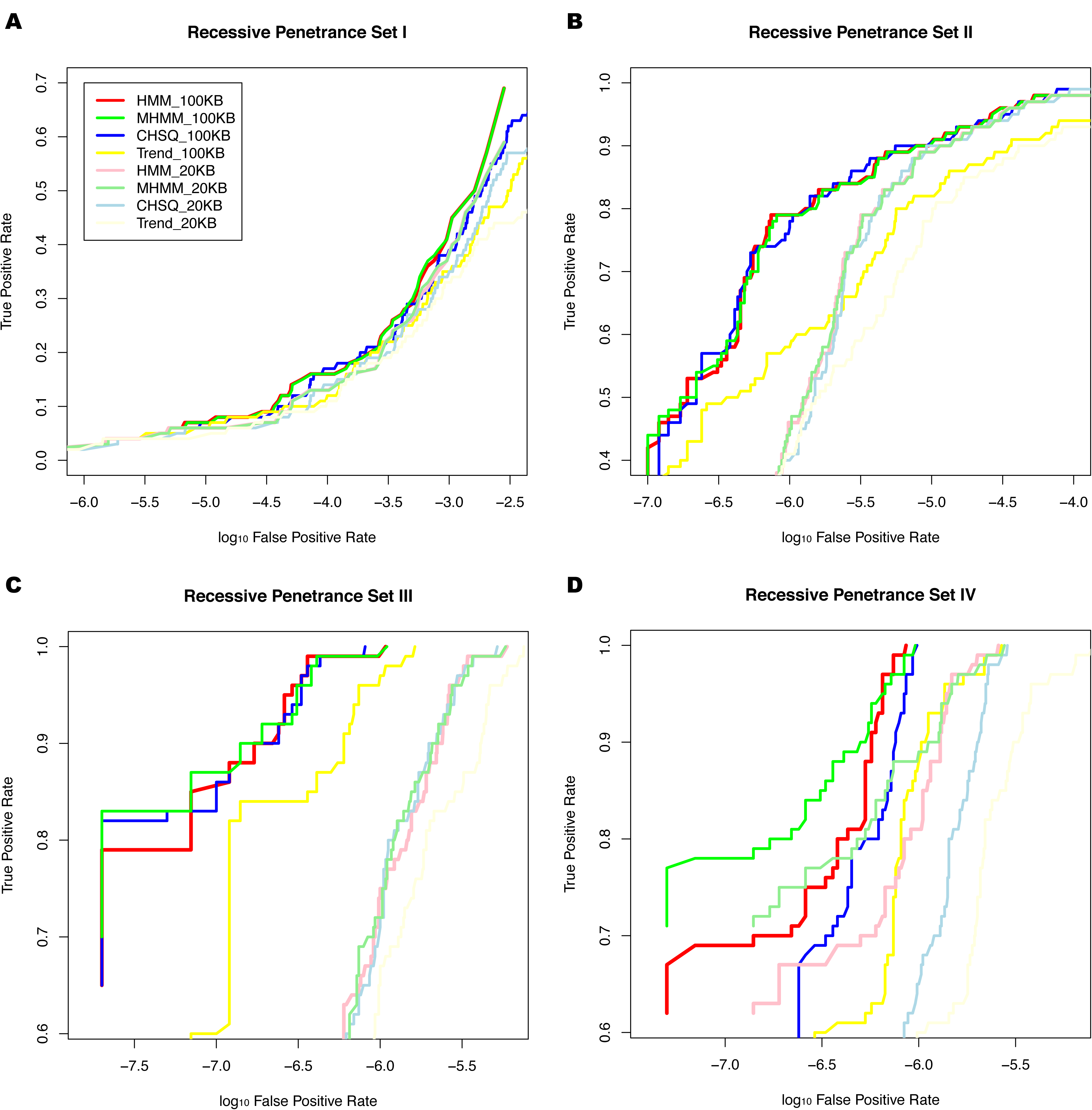
ROC curves of HMM / MHMM methods, *χ*^2^ genotypic association test, and Cochran-Armitage trend test for four recessive penetrance sets. A, B, C, and D are plots for recessive penetrance set I, II, III, and IV, respectively. The legend is the same as in Figure 2.

In the genome-wide studies done so far, the first few markers with the highest significance were of particular interest to investigators at the initial stage of association studies. In consecutive steps, these most significant markers were selected to proceed with replication analysis [6,23]. To investigate the accuracy of our methods in this manner, we also assessed the coverage of true signals among the most significant K signals at four levels of relative risks, where the rank *K* ∈ {1, 5,10,15, 20, 30, 50,100} (see Figure 4). For additive penetrance set I with *RR* = 1.25, only a couple of the true signals out of 100 are among the top 15 signals of each replication data set and the probability of covering the causal SNP is only 0.08 (even when the rank reaches 100). The HMM / MHMM shows small discrepancies compared to the *χ*^2^ test, i.e., they perform slightly better than the *χ*^2^ test at rank 50th. When *RR* increases to 1.5, the coverage probabilities of all the four methods increase substantially, almost close to 0.7 at the rank of 100. In this case, the coverage pattern of the HMM / MHMM shifts above the *χ*^2^ test. In additive penetrance set III and IV, all the methods achieve almost perfect coverage when considering the top 10 signals. The Cochran-Armitage trend test generally outperforms the other three methods at various levels of relative risk. The coverage generally decreases slightly or almost overlaps when the distance from the causal SNP decreases from 100KB to 20 KB. Under the recessive allelic effect (shown in Figure 5), the HMM / MHMM pick up the true signals more accurately than both the *χ*^2^ test and the Cochran-Armitage trend test at low or moderate levels of relative risk. With high level of *RR*, their performance are similar to each other, achieving almost 100% accuracy when the rank reaches 5. The Cochran-Armitage trend test performs poorly compared with the other three methods for small values of relative risk, consistent with previous findings.

**Figure 4.**
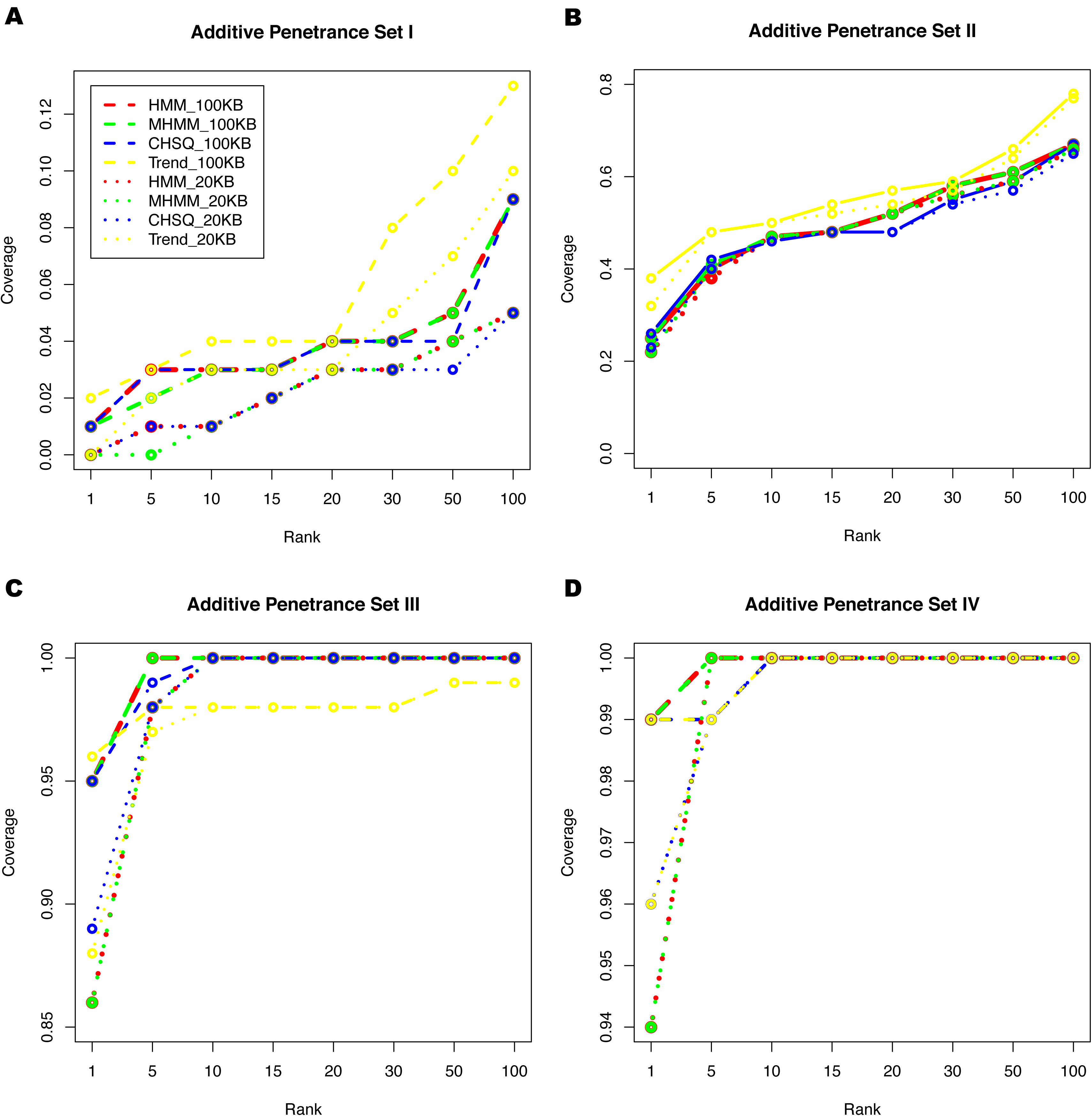
The coverage of true SNPs among the top *K* positive signals at four additive penetrance sets. *K* ∈ {1, 5,10,15, 20, 30, 50,100}. The red lines represent the ROC curves of HMM method; the green lines are for MHMM method; the blue lines are for *χ*^2^ genotypic association test; the yellow lines are for Cochran-Armitage trend test. “100KB” or “20KB” means the distance threshold from the causal variant is 100 kb or 20kb, represented by dashed and dotted lines, respectively.

**Figure 5.**
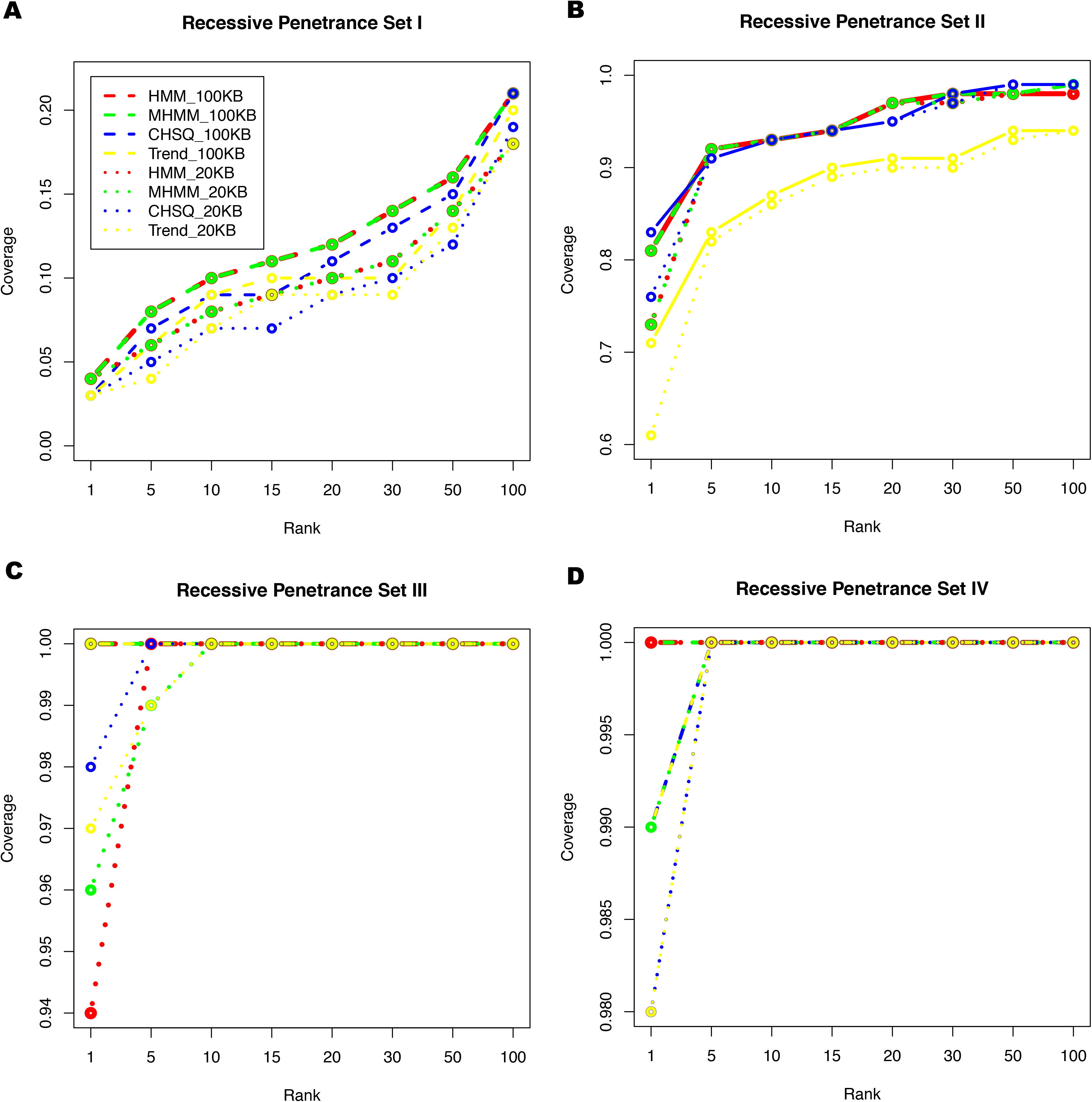
**The coverage of true SNPs among the top** *K* **positive signals at four recessive penetrance sets** *K* ∈ {1, 5,10,15, 20, 30, 50,100}. The legend is the same as in Figure 4.

### Application to WTCCC Data

To demonstrate a concrete application of HMM / MHMM methods, we have applied this approach to the large amount of data from [6] generated by the Affymetrix GeneChip 500K Array Set, containing about 3000 shared controls and about 2000 cases for each of the seven human common diseases, including bipolar disorder, coronary artery disease (CAD), Crohns disease, hypertension, rheumatoid arthritis, type 1 diabetes, and type 2 diabetes. Our HMM / MHMM model is such a flexible framework that it can be applied to the summarized data downloaded from this study directly. The prior probability of associated state is chosen as 10^−6^, the measure of LD is 1/15kb as before and the non-centrality parameter is 50. Figure 6 illustrated the performance of HMM / MHMM methods applied to the case-control data set for coronary artery disease. Our methods also detect the strongest signal occurred at the SNP rs1333049 with posterior probability close to 1, which has been confirmed by multiple studies in various populations [24-27] to confer CAD risk to patients. The results of our method applied to other disease data sets are shown in Supplemental Materials. Generally, the background noise is minimized after using our methods, compared with the cases using either the *χ*^2^ genotypic association test or the Cochran-Armitage trend test.

**Figure 6.**
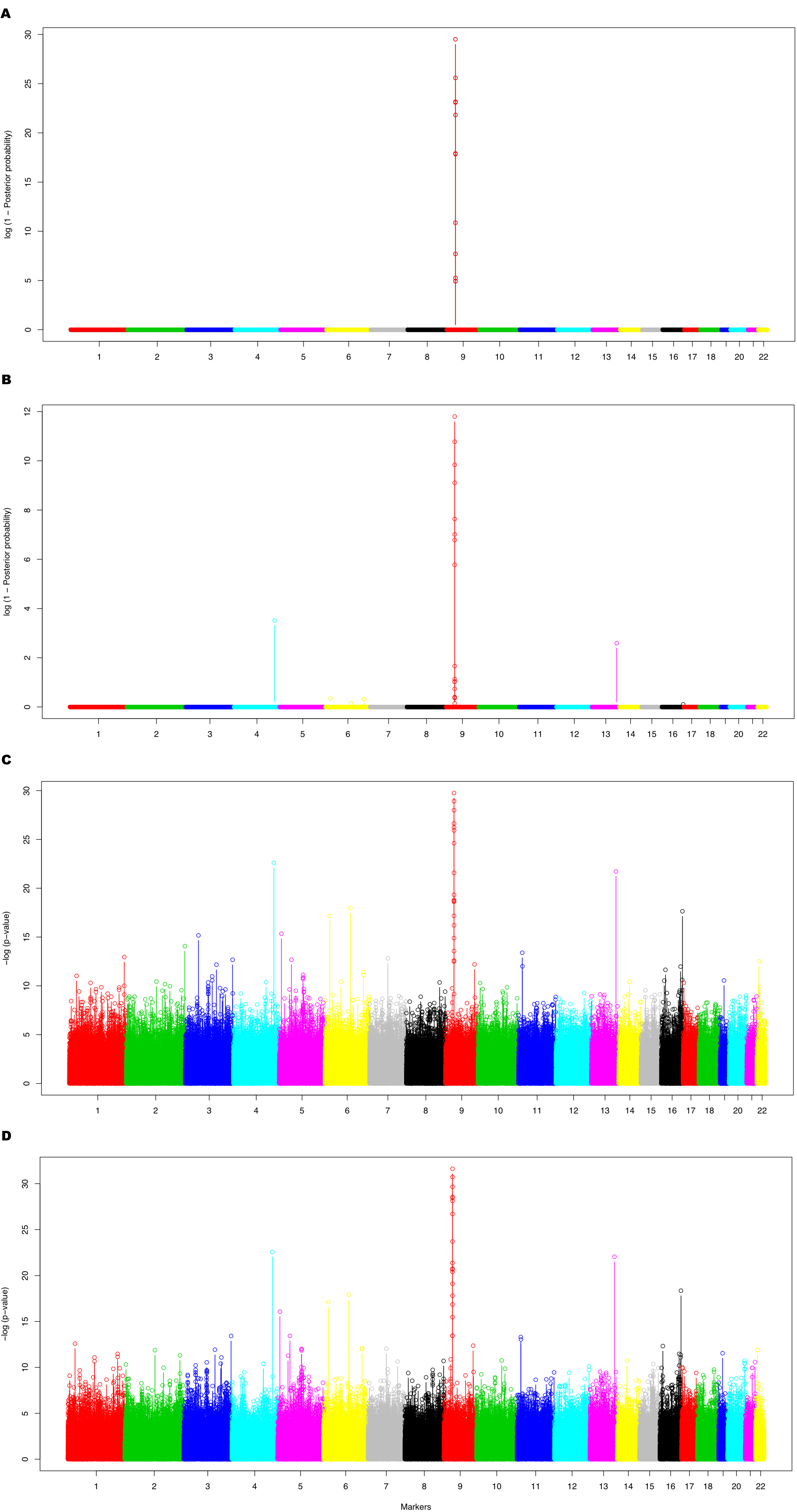
Association of markers across the whole genome from the WTCCC data set for coronary artery disease produced by the HMM method (A), the MHMM method (B), *χ*^2^ genotypic association test (C) and Cochran-Armitage trend test (D). The y-axis in subfigures A and B represents log(1 Posterior probability), where “Posterior probability” is the posterior probability of a marker in the associated state obtained from the HMM / MHMM method. The y-axis in subfigures C and D represents – log(p-value). The tick-marks on the x-axis represent the chromosome where markers are located. Different colors separate markers on different chromosome.

## Discussion

In this paper, we present an original and efficient method for genome-wide association mapping. Our method utilizes the succinct and flexible framework of the hidden Markov model and its derivate the Markov hidden Markov model to account for linkage disequilibrium among markers, which effectively removes background noises. It then estimates posterior probabilities of markers associated with phenotypes to make inference in mapping risk genetic variants, naturally avoiding the multiple comparison issue. Extensive simulations illustrate that our method outperforms most popular methods used in GWAS, including the *χ*^2^ genotypic association test and the Cochran-Armitage trend test, when the allelic effect is not exactly additive or we are lack of prior information about the allelic effect of the causal variant.

Our HMM / MHMM approaches performs superiorly to other methods in that it utilizes the linkage disequilibrium among markers to clear out almost all the background noises. In this way, even weak signals are magnified substantially and become easy to detect, as long as the effects of signals are above certain detectable threshold of our method. If below the detectable threshold, our method will classify them as background noises and the signals are weakened. Due to its well control of Type I error, our method attains higher power than most other methods at fixed false positive rates when the allelic effect is non-additive and the strength of signals is low to moderate. When the signals are strong and detectable for most methods, the performance of our method is similar to that of *χ*^2^ genotypic association test and the Cochran-Armitage trend test. We also observed in our simulations that the Cochran-Armitage trend test outperforms other methods with the additive SNP effect, but performs the worst when the allelic effect is recessive. According to Sasieni [9], the Cochran-Armitage trend test using the number of one allele as weights is locally most powerful only when the allele effect is exactly additive (i.e., if the homozygous odds ratio is the square of the heterozygous one), which explains the observed paradox. This result implies that the Cochran-Armitage trend test should be used only when the allelic effect of the causal SNP is known to be additive; otherwise, the HMM / MHMM is a more appropriate approach to detect associations.

It is important to note that the power of our method depends on multiple factors at fixed false positive rates. First of all, it is intimately related to the pattern of the linkage disequilibrium around the causal variant. If there are a few flanking markers tightly linked to the causal one, the potential of detecting it is relatively high as long as this locus has at least moderate trait effect. Otherwise, it is rather difficult to uncover its location as the effect of this single signal might be obscured when clearing out background noises. Moreover, the transition matrix used in the HMM / MHMM setting greatly influence its performance. In particular, the *β* parameter is key since it determines that how frequently the Markov chain is expected to enter the associated state. If we employ the widely-used exponentially-decay transition matrix [18,19,28], ideally the stationary distribution of an Markov chain is governed by the prior probability of being in the association state [19]. However, when the distance between some consecutive markers becomes as small as 100 bp, the transition matrix converges to an identity matrix with two states. This leads to inflated false positives wherever marker sets are dense. We suggest that when markers are equally spaced with distances between two consecutive loci greater than 1KB, it is appropriate to apply the original exponentially-decay transition matrix; otherwise, to avoid the above problem, we use the matrix presented in Method section, whose stationary distribution is determined by both the prior probability and the amount of exponential decay in LD.

In addition, the parameters in the models influence the estimation of posterior probabilities as well as power. The prior probability of state A has a slim influence on the accuracy of detecting causal SNPs as from extra simulation, the pattern of ROC curves of our method shifts slightly to the right even when the value of *β* is increased by four magnitudes (results not shown). And the ranking of markers in terms of posterior probabilities is not affected by the value of the prior probability either. Another variable that influences the performance is the LD decay parameter λ. Here we have used the value of 1/15kb to reflect average decay of LD in Europeans (approximately 50 kb) (see Figure 1). Simulations indicate that our model is fairly robust to the misspecification of this parameter (data not shown). The non-centrality parameter has a subtle effect on the power as long as it is larger than the non-centrality estimators of markers with the most skewed contingency tables. The correlation coefficient in the joint *χ*^2^ distribution also plays an important role in estimation accuracy of MHMM. Current practice is to estimate the correlation coefficient using exponential decay with distance between consecutive markers, which is a very rough approximation. A better way to estimate it is to obtain an estimator from the posterior distribution of the correlation coefficient based on prior information and the sample correlation coefficient [29]. Although these parameters do influence the power, we do not encourage the use of Markov chain Monte Carlo methods to integrate out the uncertainty of these parameters given empirical information about them, since it is not worthwhile to sacrifice the speed and efficiency of these models, especially with large data sets.

It is important to note that our current implementation ignores population structure in the data, which may be an important confounding variable in some applications. Our simulated data are constituted solely of Europeans extracted from the GSK project. In spite of the fact that this dataset contains individuals simply of the same ethnicity, there is inevitably some population differentiation within the data set. Several projects [6, 8, 10, 23] also chose individuals from identical ethnic backgrounds to avoid a high degree of population subdivision. However, false positives caused by population stratification cannot be totally eliminated. Recently, a few methods have been proposed to correct for substructure, such as genomic control [30-32], structure association [33-35], unified mixed model [36] and Eigenstrat [37]. Compared with other methods, the Eigenstrat approach is computationally efficient and achieves the lowest error rates in correcting for population structure. Additionally, it can be easily incorporated into our HMM / MHMM. Eigenstrat works by regressing both the phenotypic and genotypic values on the individual ancestry coefficients obtained from principal component decomposition. Such covariates can be included within the HMM / MHMM analysis via the use of standard residuals in linear regression.

A further step that might make the inference more reliable is to estimate penetrances of potential markers discovered by the HMM / MHMM models using the method proposed by Zollner et al. [38]. If the relative risk obtained from penetrance estimation is around 1.5 or less, we recommend further replication studies on the top 100 signals or more; if the relative risk is around 1.75 or above, we suggest only the top 10 SNPs need to be investigated in replication studies as the coverage of the true SNP among the top 10 signals is almost 100% at this level of relative risk (see Figure 4 and 5).

Our HMM / MHMM method is rather simple and effective for detecting disease-related polymorphisms. It achieves higher power at fixed Type I error rates, than most popular statistical tests, due in part to the utilization of the correlation information among markers and the correction for background linkage disequilibrium among dense markers. It is a flexible framework that can accommodate various factors influencing power, such as population stratification and other potential covariates. We anticipate this method can be practically applied to many association mapping projects across a wide spectrum of disease phenotypes.

## Methods

We assume a sample of *N* individuals have been genotyped at *L* linked SNP loci. Let state “*NA*” represent that a marker of interest is not associated to a phenotype, and state “*A*” means the marker is associated to a phenotype. According to Tang et al. [19], the three major elements of the HMM / MHMM frameworks are the prior distribution of the two states, the transition probability matrix, and the emission probabilities. We used β to represent the prior probability of being in the associated state. Since the number of disease-predisposing regions is very limited for most applications, *β*’s value is small and close to zero. (In practice, we can think of *β* as a measure of the investigators cost of follow-up. Large values of *β* will produce more associated regions, and small values of *β* will reduce the number of potentially associated regions for follow-up). To model the mosaic configuration of chromosomes, we employ a Poisson process similar to those used in admixture inference [15-17, 19]. The breakpoints between strong LD patches along the chromosomes are randomly picked at the rate λ per kb, where λ reflects the extent of linkage disequilibrium decay with distance across the genome. At stationary, this process produces signals of association with frequency of the prior probability of state *A*. However, when using GWAS, markers cover most of the genome so densely that this transition matrix is prone to generate false positive signals as it approximates the identity matrix (we provide a detailed explanation in Discussion section). To mitigate this problem, we use a modified transition matrix:

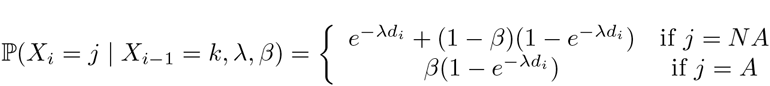

where *d_i_* is the physical distance between marker *i* and marker *i* - 1. In this transition matrix, the stationary probability of staying in state *A* is approximately 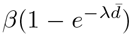, influenced by both the prior probability and the LD decay measure.

The observation at marker *i*, denoted by *Y_i_*, is Pearson’s *χ*^2^ goodness-of-fit test statistic based on the allelic or genotypic contingency table. Considering the case-control genotypic contingency table, let {*n_ij_*: *i* = 1 or 2; *j* = 1, 2, or 3} denote the observed frequencies of genotypes among cases or controls, i.e. values in the cells of the contingency table. Let {*p_ij_*: *i* = 1 or 2; *j* = 1, 2, or 3} represent the probabilities of these cells under the null hypothesis of no association. The *χ*^2^ test statistic is just calculated through the formula 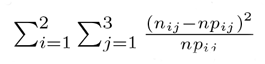, where n is just the sum of all *n_ij_*’*s*. This procedure of generating observation *Y_i_* can be easily extended to the ordinal phenotype case.

The emission probability of this statistic, under the state *NA*, i.e., the null hypothesis that marker i is not associated with the phenotype, is approximated by the density of the central chi-square distribution with two degrees of freedom. Under the alternative state that marker *i* is associated with the phenotype, the asymptotic distribution of *Y_i_* is the non-central chi-square distribution with five degrees of freedom and the non-centrality parameter *κ* representing its skew parameter (its probability density is 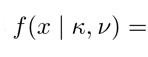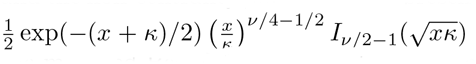, where *x* ~ *nc* – *χ*^2^ with *ν* degrees of freedom and *I_ν_*(*z*) is the modified Bessel function of the first kind). From the practical perspective, it is ideal to attain high probability under alternative hypothesis and low probability under null hypothesis for a skewed binary contingency table, which implies the ability to distinguish associated signals from background noises. Therefore, we recommend choosing a non-centrality parameter large enough to bound most skew cases in association mapping.

The Markov hidden Markov model we consider models extra correlation among the observations of consecutive markers when both markers are in the same state. This is a major difference between MHMM and HMM. In terms of likelihood computation, this amounts to calculating the emission probability of marker *i* as the conditional probability of its observation given the previous observation and their congruent hidden states, defined as

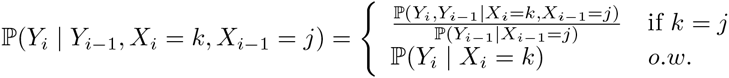

where P(*Y_i_*, *Y*_*i*-1_| *X_i_*, *X*_*i*-1_) represents the joint distribution of observations at two consecutive loci provided that their hidden states are identical. The joint distributions is a bivariate central chi-square distribution with two degrees of freedom under the non-associated state 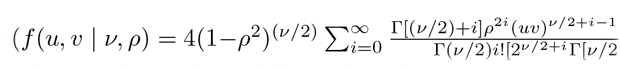 where *u*, *υ* ~ *χ*^2^ with *ν* degrees of freedom). [39] Under the associated state, the statistics follow a bivariate non-central chi-square distribution with five degrees of freedom and the non-centrality parameter and the correlation parameter 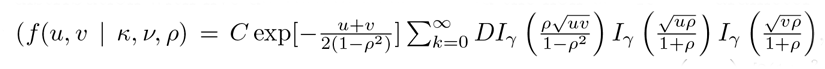, where *u*, *υ* ~ *nc* – *χ*^2^ with *ν* degrees of freedom and the non-centrality parameter *κ*, 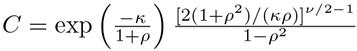,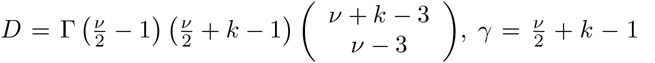 and *I_γ_* (*z*) is the modified Bessel function of the first kind [40]). We have implemented the above models in an ANSI C program, *HMM*_*Map*, which is available upon request.

When the phenotypes are continuous, the observed statistic at marker *i*, also denoted by *Y_i_*, is F test statistic after the linear regression of the trait values on the genotypic values of marker *i*, *X_i_*, i.e., suppose *Y_i_* = *βX_i_* + *∊*, where *β* is the regression parameter and *∊* represents random error, then the F statistic is 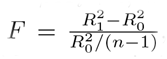, where 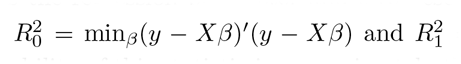 and 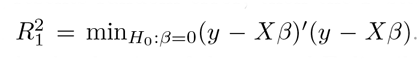. For HMM, the emission probability of this statistic is approximately the density of the central F distribution *F*(1, *n* - 1) under the null hypothesis that marker *i* is not associated with the phenotype. Whereas under the associated state that marker *i* is associated with the phenotype, the asymptotic distribution of *Y_i_* is the non-central F distribution *F*(1,*n* – 1, *κ*), where *κ* is the non-centrality parameter representing its asymmetric level. Under Markov hidden Markov model, the only difference is that we need to compute the joint distribution of observations at two consecutive markers given their hidden states are the same, which follows bivariate central F distribution under the non-associated state or follows bivariate non-central F distribution with the non-centrality parameter and the correlation parameter.

### Inference

For the HMM model, the likelihood function can be computed directly using the standard forward algorithm. Likewise, the modified forward algorithm can be used to calculate the likelihood under the MHMM model [19]. Three parameters need to be specified for both: the prior probability of state *A*, *β*, the measure of LD, *λ*, and the non-centrality parameter, *κ*. Although it is straightforward to integrate out uncertainty in these parameters via Markov chain Monte Carlo (MCMC) algorithm, such an approach can be quite time consuming. Since efficiency is critical in genome-wide studies, we have used heuristic estimates of these parameters in our likelihood calculations. The posterior probability of a marker being in the associated state is calculated using the forward-backward algorithms for the HMM and the semi-forward-backward algorithm developed in [19] for MHMM setting. This statistic is a natural choice for an association measure between marker genotype and phenotype. For the purpose of method comparisons, we also applied the *χ*^2^ test for genotype association and Cochran-Armitage trend test on each marker.

### Data Simulation

We randomly selected 1,000 Europeans with 22 autosomes from the data obtained from the GSK-POPRES project. (This data set contains Affymetrix 500K genotype data on 3845 samples from around the world, consisting of 443,434 SNPs.) After removing loci without polymorphism, 416,331 SNPs were retained. For each of 100 replications generated based on this data set, the causal SNP was randomly picked across the genome, among SNPs with minor allele frequencies (MAF) above 0.3. (In the sampling design of case-control studies, the number of cases is usually on the comparable magnitude as controls. Thus it is necessary to choose an SNP with large MAF). Phenotypes were then generated according to the SNP genotypes and penetrances defined as the probability of an individual having the disease, given his or her genotype at this locus. Here we considered both additive and recessive allelic models for generation of phenotypes (dominant model is neglected here as it is similar to recessive model). For each allelic model, four penetrance sets in Table ?? were chosen to cover the range of moderate to low relative risks (*RR* ∈ {1.25,1.5,1.75, 2.0}).

## Acknowledgments

This work is funded by NIH(NIGMS) 1R01GM83606 to Carlos D. Bustamante, Sara Tishkoff, Martin T. Wells, and Jason Mezey.

## References

1. International HapMap Consortium (2003) The International HapMap project. Nature 426: 789–96.

2. International HapMap Consortium (2005) A haplotype map of the human genome. Nature 437: 1299–320.

3. International HapMap Consortium (2007) A second generation human haplotype map of over 3.1 million snps. Nature 449: 851–61.

4. Hirschhorn J, Daly M (2005) Genome-wide association studies for common diseases and complex traits. Nat Rev Genet 6: 95–108.

5. Wang W, Barratt B, Clayton D, Todd J (2005) Genome-wide association studies: theoretical and practical concerns. Nat Rev Genet 6: 109–18.

6. Wellcome Trust Case Control Consortium (2007) Genome-wide association study of 14,000 cases of seven common diseases and 3,000 shared controls. Nature 447: 661–78.

7. Abecasis G, Yashar B, Zhao Y, Ghiasvand N, Zareparsi S, et al. (2004) Age-related macular degeneration: a high-resolution genome scan for susceptibility loci in a population enriched for late-stage disease. Am J Hum Genet 74: 482–94.

8. Klein R, Zeiss C, Chew E, Tsai J, Sackler R, et al. (2005) Complement factor H polymorphism in age-related macular degeneration. Science 308: 385–9.

9. Sasieni PD (1997) From genotypes to genes: doubling the sample size. Biometrics 53: 1253–1261.

10. Sladek R, Rocheleau G, Rung J, Dina C, Shen L, et al. (2007) A genome-wide association study identifies novel risk loci for type 2 diabetes. Nature 445: 881–5.

11. Benjamini Y, Hochberg Y (1995) Controlling the false discovery rate: a practical and powerful approach to multiple testing. J R Statist Soc B 57: 289–300.

12. Storey J (2002) A direct approach to false discovery rates. J R Statist Soc B 64, part3: 479–98.

13. Storey J, Tibshirani R (2003) Statistical significance for genomewide studies. Proc Natl Acad Sci 100: 9440–5.

14. Yang Q, Cui J, Chazaro I, Cupples L, Demissie S (2005) Power and type I error rate of false discovery rate approaches in genome-wide association studies. BMC Genet 6 Suppl. 1: S134.

15. Hoggart C, Parra E, Shriver M, Bonilla C, Kittles R, et al. (2003) Control of confounding of genetic associations in stratified populations. Am J Hum Genet 72: 1492–504.

16. Patterson N, Hattangadi N, Lane B, Lohmueller K, Hafler D, et al. (2004) Methods for high-density admixture mapping of disease gene. Am J Hum Genet 74: 979–1000.

17. Montana G, Pritchard J (2004) Statistical tests for admixture mapping with case-control and cases-only data. Am J Hum Genet 75: 771–89.

18. Falush D, Stephens M, Pritchard J (2003) Inference of population structure using multilocus genotype data: linked loci and correlated allele frequencies. Genetics 164: 1567–87.

19. Tang H, Coram M, Wang P, Zhu X, Risch N (2006) Reconstructing genetic ancestry blocks in admixed individuals. Am J Hum Genet 80: 353–60.

20. Auton A, Bryc K, Boyko A, Lohmueller K, Novembre J, et al. (2009) Global distribution of genomic diversity underscores rich complex history of continental human populations. Genome Res 19: 795-803.

21. Metz C (1978) Basic principles of roc analysis. Semin Nucl Med 8: 283–298.

22. Zhang M, Montooth KL, Wells MT, Clark AG, Zhang D (2005) Mapping multiple quantitative traite loci by Bayesian classification. Genetics 169: 2305–2318.

23. Karlsson E, Baranowska I, Wade C, Salmon H, Hc N, et al. (2007) Efficient mapping of mendelian traits in dogs through genome-wide association. Nat Genet 39: 1321–8.

24. Samani N, Erdmann J, Hall A, Hengstenberg C, Mangino M, et al. (2007) Genomewide association analysis of coronary artery disease. N Engl J Med 357: 443–453.

25. Bown M, Braund P, Thompson J, London N, Samani N, et al. (2008) Association between the coronary artery disease risk locus on chromosome 9p21.3 and abdominal aortic aneurysm. Circulation: Cardiovascular Genetics 1: 39–42.

26. Hinohara K, Nakajima T, Takahashi M, Hohda S, Sasaoka T, et al. (2008) Replication of the association between a chromosome 9p21 polymorphism and coronary artery disease in japanese and korean populations. Journal of Human Genetics 53: 357–359.

27. Schunkert H, Gotz A, Braund P, McGinnis R, Tregouet D, et al. (2008) Repeated replication and a prospective meta-analysis of the association between chromosome 9p21.3 and coronary artery disease. Circulation 117: 1675–1684.

28. Morris A, Whittaker J, Balding D (2000) Bayesian fine-scale mapping of disease loci, by hidden Markov models. Am J Hum Genet 67: 155–69.

29. Schisterman E, Moysich K, England LJ, Rao M (2003) Estimation of the correlation coefficient using the Bayesian Approach and its applications for epidemiologic research. BMC Medical Research Methodology 3.

30. Reich D, Goldstein D (2001) Detecting association in a case-control study while correcting for population stratification. Genet Epidemiol 20: 4–16.

31. Devlin B, Roeder K (1999) Genomic control for association studies. Biometrics 55: 997–1004.

32. Devlin B, Bacanu S, Roeder K (2004) Genomic control to the extreme. Nat Genet 36: 1129–30.

33. Pritchard J, Stephens M, Rosenberg N, Donnelly P (2000) Association mapping in structured populations. Am J Hum Genet 67: 170–81.

34. Satten G, Flanders W, Yang Q (2001) Accounting for unmeasured population substructure in case-control studies of genetic association using a novel latent-class model. Am J Hum Genet 68: 466–77.

35. Setakis E, Stirnadel H, Balding D (2006) Logistic regression protects against population structure in genetic association studies. Genome Res 16: 290–6.

36. Yu J, Pressoir G, Briggs W, Vroh Bi I, Yamasaki M, et al. (2006) A unified mixed-model method for association mapping that accounts for multiple levels of relatedness. Nat Genet 38: 203–8.

37. Price A, Patterson N, Plenge R, Weinblatt M, Shadick N, et al. (2006) Principal components analysis corrects for stratification in genome-wide association studies. Nat Genet 38: 904–9.

38. Zollner S, Pritchard J (2007) Overcoming the winner’s curse: estimating penetrance parameters from case-control data. Am J Hum Genet 80: 605–15.

39. Krishnaiah P, Hagis P, Steinberg L (1963) A note on the bivariate chi distribution. SIAM Review 5: 140–4.

40. Krishnan M (1967) The noncentral bivariate chi distribution. SIAM Review 9: 708–14.

41. Ardlie K, Kruglyak L, Seielstad M (2002) Patterns of linkage disequilibrium in the human genome. Nat Rev Genet 3: 299–309.

